# Viral infection causes sex-specific changes in fruit fly social aggregation behaviour

**DOI:** 10.1101/630913

**Authors:** Jonathon A. Siva-Jothy, Pedro F. Vale

## Abstract

Host behavioural changes following infection are common and could be important determinants of host behavioural competence to transmit pathogens. Identifying potential sources of variation in sickness behaviours is therefore central to our understanding of disease transmission. Here, we test how group social aggregation and individual locomotor activity vary between different genotypes of male and female fruit flies (*Drosophila melanogaster*) following septic infection with Drosophila C Virus. We find genetic-based variation in both locomotor activity and social aggregation but we did not detect an effect of DCV infection on fly activity or sleep patterns within the initial days following infection. However, DCV infection caused sex-specific effects on social aggregation, as male flies in most genetic backgrounds increased the distance to their nearest neighbour when infected. We discuss possible causes for these differences in the context of individual variation in immunity and their potential consequences for disease transmission.

## Introduction

Infection-induced changes to host physiology, and immunity in particular, following infection are well known, but it is equally striking that many animals experience similar behavioural changes following infection [1,2]. Common behavioural responses to infection include eating and moving less, as well as foregoing social and sexual interactions [1,3–5]. Whether these behavioural changes in response to infection are evolved host responses, parasite manipulations, or a coincidental by-product of infection[6,7], they have potentially important consequences for disease transmission [8]. This is particualry clear for behaviours such as individual locomotor activity or group social aggregation, which will directly determine how frequently susceptible and infected individuals interact. Assessing how host behaviours that influence contact rates might change following infection is therefore central to understanding the spread of infectious disease.

The extent to which host behaviours are modified during infection is likely to depend on genetic and environmental factors. Even in the absence of infection, individuals of some genetic backgrounds are more likely to aggregate than others [9,10], while males and females in a broad range of host species often exhibit distinct behavioural profiles [11,12]. How these different sources of variation influence infection-induced behavioural changes is relatively understudied [8]. Measuring how males and females of different genetic backgrounds modify their behaviour during infection may highlight groups of individuals with higher contact rates and offer insight into the potential causes of heterogeneity in pathogen spread.

Testing if locomotor and aggregation behaviours change following infection, and if these changes differ between genetic backgrounds, is not straightforward for most host species. It requires knowledge of how individuals within a population differ in their genetic backgrounds and the ability to expose many individuals of the same background to infection in controlled conditions, while comparing their behavioural responses to infection with individuals of the same background that do not experience infection. For many animal species, and certainly in human populations, this type of intervention is either extremely challenging or not feasible. One alternative is to leverage the tools offered by model systems. *Drosophila melanogaster*, for example, has been widely used as a model system for behavioural genetics [13,14], and used specifically to study social aggregation and locomotor activity [9,15,16]. Further, *D. melanogaster* is a powerful model of immunity in response to a range of bacterial and viral pathogens [17]. Previous work has shown that *D. melanogaster* exhibits a range of behavioural changes following *Drosophila* C Virus (DCV) infection, including pathogen avoidance during oviposition [18], and foraging [19]. Here, we test whether DCV infection changes social aggregation and locomotor activity in *D. melanogaster*, and if these effects vary with genetic background and sex.

## Materials & Methods

### Flies and Rearing Conditions

We used males and females from 10 lines sourced from the Drosophila Genetic Resource Panel (DGRP) [20], and are among the most and least susceptible genetic backgrounds to systemic Drosophila C Virus infection [21]. Flies were reared in plastic vials on a standard diet of Lewis medium at 18±1°C with a 12 hour light:dark cycle with stocks tipped into new vials every 14 days. One month before the experiment, flies were transferred to incubators and maintained at 25±1°C with a 12 hour light:dark cycle at low density (∼10 flies per vial) for two generations.

### Virus Culture and Infection

The Drosophila C Virus (DCV) isolate was originally isolated in Charolles, France [22] and the stock used in this experiment was grown in Schneider Drosophila Line 2 (DL2) as previously described [23] diluted ten-fold (10^8^ infectious units per ml) in TRIS-HCl solution (pH=7.3), aliquoted and frozen at −70°C until required. To infect with DCV, 3-5 day old flies were pricked in the pleural suture with a 0.15mm diameter pin, bent at 90° ∼0.5mm from the tip, dipped in DCV (or TRIS-HCl for controls), as described previously [24,25].

### Measuring Drosophila activity

The activity of single flies was measured during 4 continuous days using a Drosophila Activity Monitor (DAM2 System, TRIKinetics), in an incubator maintained at 25°C in a 12:12 light:dark cycle [15]. In total we measured the activity of 872 flies, with n=18-27 flies for each combination of sex and genetic background (Table S3). Raw activity data was processed using the DAM System File Scan Software [15], and the resulting data was manipulated using Microsoft Excel. We analysed fly activity data using three metrics: total locomotor activity, proportion of time spent asleep and the average activity when awake, as described previously [23]. A more detailed description of the experimental setup and analysis can be found in electronic supplementary material.

### Measuring Drosophila social aggregation

Social aggregation was measured in a separate experiment, by calculating the nearest neighbour distance (NND) between each individual fly within a 12-fly group contained within a Petri dish [10,16,26]. The experiment was conducted over five experimental blocks, each carried out over a single day, where each genetic background, sex and infection treatment was measured. The NND was calculated by image analysis of pictures recorded of each group using the ‘NND’ package in ImageJ [27]. In total, we measured social aggregation in 580 groups of flies, with n=14-16 replicate groups of 12 flies for each genetic background, sex and infection status combination. A more detailed description of the experimental setup and analysis can be found in electronic supplementary material.

### Statistical Analysis

We tested if differences in locomotor activity and social aggregation could be attributed to fly genetic background or sex. Data from both experiments were analysed using very similar Generalized Linear Models (GLMs). Models used a full factorial 3-way interaction between infection status (control /infected), sex (male / female) and DGRP line (10 lines), all modelled as fixed effects. Analysis of social aggregation used a model listing only the median nearest neighbour distance of each dish as its response variable. To assess locomotor activity, we analysed 3 response variables in separate GLMs (total activity, proportion of time asleep, awake activity), adjusting the significance threshold to 0.01667 using Bonferroni correction to account for multiple-testing. All statistical analyses and graphics were carried out and produced in R 3.3.0 using the *ggplot2* (Wickham, 2016) and *lme4* (Bates et al., 2015) packages.

## Results

### Social aggregation

We found a significant effect of genetic background in the median nearest neighbour distances (NND), suggesting that there is significant genetic variation in this measure of social aggregation (Figure 1; Table 1). We found no evidence of sexual dimorphism in social aggregation across multiple genetic backgrounds, with no significant interaction between sex and genetic background. However, we observed that while female aggregation was not affected by infection, infected males aggregated further apart from each other compared to uninfected males (Figure 1; Table 1). This increase in the NND following infection was generally observed in males, regardless their genetic background (Figure 1; Table 1). We also detected an expected sexual dimorphism in body size, as female *D. melanogaster* are typically larger than males (Figure S1). Incorporating this size difference into measures of social aggregation, by measuring body lengths between individuals did not alter the results qualitatively (Figure S2).

**Table 1.** Model outputs for statistical tests performed on social aggregation, testing the causes of variation in sociality in males and females of 10 *D. melanogaster* genetic backgrounds. Significant predictors are marked with asterisks (p<0.05=*, p<0.01=** and p<0.001=***).

| Response | Predictor | Df | F | p |
| --- | --- | --- | --- | --- |
| Median NND | Genetic Background | 9 | 5.0249 | <0.0001 |
|  | Sex | 1 | 2.7870 | 0.096 |
|  | Infection | 1 | 21.1301 | <0.0001 |
|  | Genetic Background * Sex | 9 | 1.4112 | 0.18 |
|  | Genetic Background * Infection | 9 | 0.9654 | 0.47 |
|  | Sex * Infection | 1 | 19.6600 | <0.0001 |
|  | Genetic Background * Sex * Infection | 9 | 1.6729 | 0.92 |

**Figure 1.**
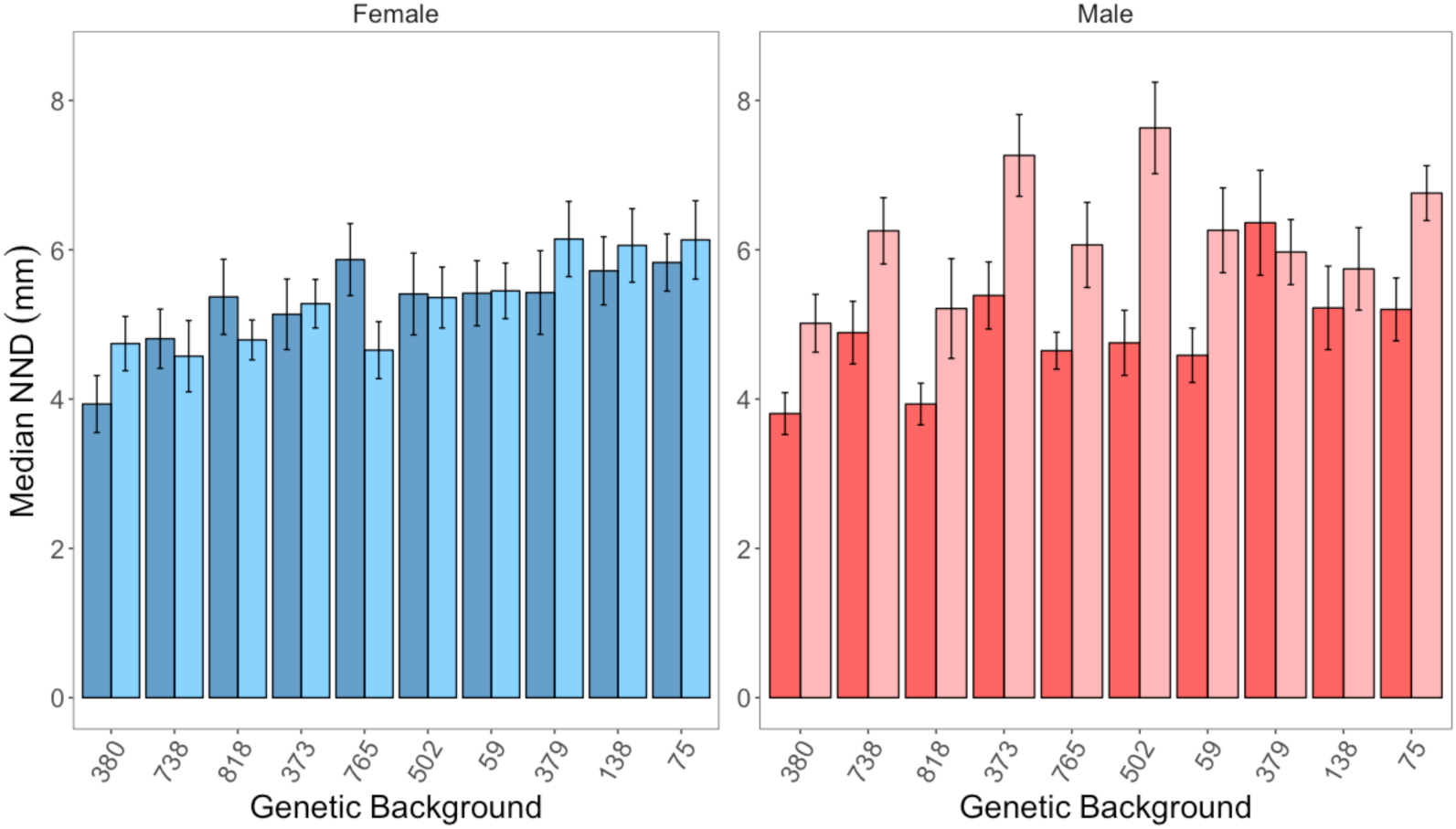
Mean±SE median nearest neighbour distance (NND) in millimetres (mm) of adult flies placed in Petri dishes for at least 30 minutes until settled. (a) Uninfected female-only arenas shown in blue, and infected female-only arenas in pale blue. (b) Uninfected male-only arenas are shown in red, and infected male-only arenas in pink. The x-axis of both panels is ordered from the lowest to highest mean median NND of female flies of a single genetic background.

### Locomotor activity

All three parameters of total locomotor activity, the proportion of time spent asleep and the average activity when awake, were affected by a combination of sex and genetic background (Figures 2 and S3; Table 2). However, there was no detectable difference in how much infected and healthy flies moved or slept, and hence no evidence that infection impacted on any parameter of fly locomotor activity (Figures 2 and S3; Table 2).

**Table 2.** Model outputs for statistical tests performed on host activity data, testing the causes of variation in locomotor activity, sleep patterns and average awake activity in males and females of 10 D. melanogaster genetic backgrounds. Significance thresholds are corrected for multiple testing using Bonferroni correction, with significant predictors are marked with asterisks (p<0.01667=*, p<0.001=** and p<0.0001=***).

| Response | Predictor | Df | F | p |
| --- | --- | --- | --- | --- |
| Total Activity | Genetic Background | 9 | 14.83 | <0.0001*** |
|  | Sex | 1 | 1.537 | 0.21 |
|  | Infection | 1 | 0.117 | 0.73 |
|  | Genetic Background * Sex | 9 | 3.0485 | 0.0013* |
|  | Genetic Background * Infection | 9 | 1.4125 | 0.18 |
|  | Sex * Infection | 1 | 3.9707 | 0.047 |
|  | Genetic Background * Sex * Infection | 9 | 1.9471 | 0.043 |
| Proportion of Time Spent Asleep | Genetic Background | 9 | 25.1759 | <0.0001*** |
|  | Sex | 1 | 77.9823 | <0.0001*** |
|  | Infection | 1 | 0.6939 | 0.41 |
|  | Genetic Background * Sex | 9 | 3.444 | <0.001** |
|  | Genetic Background * Infection | 9 | 0.8021 | 0.61 |
|  | Sex * Infection | 1 | 0.7513 | 0.39 |
|  | Genetic Background * Sex * Infection | 9 | 1.4612 | 0.16 |
| Awake Activity | Genetic Background | 9 | 8.1673 | <0.0001*** |
|  | Sex | 1 | 0.6641 | 0.42 |
|  | Infection | 1 | 0.0008 | 0.98 |
|  | Genetic Background * Sex | 9 | 5.2153 | <0.0001*** |
|  | Genetic Background * Infection | 9 | 0.8716 | 0.55 |
|  | Sex * Infection | 1 | 0.8430 | 0.36 |
|  | Genetic Background * Sex * Infection | 9 | 1.2998 | 0.23 |

**Figure 2.**
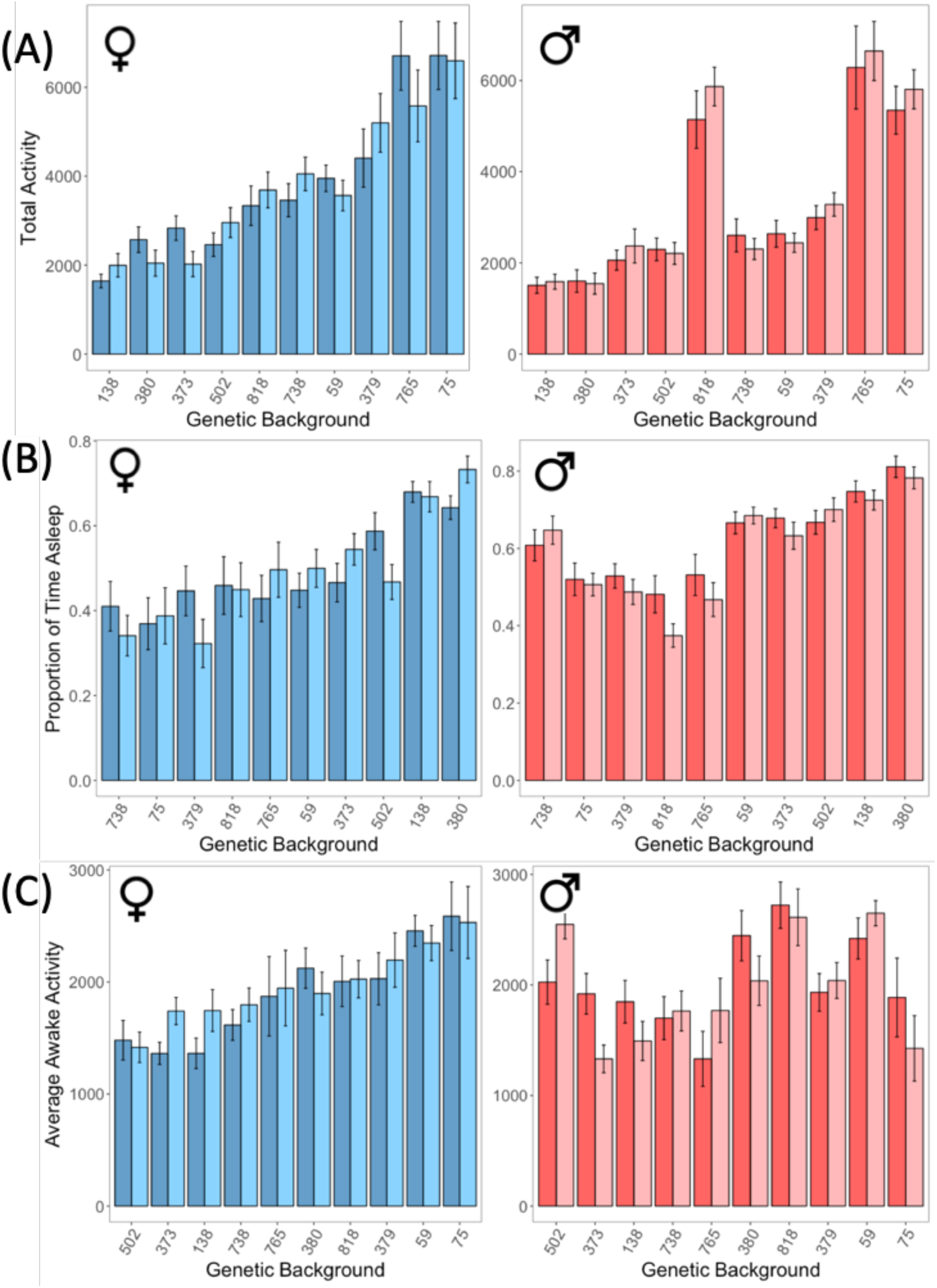
Mean±SE (A) total locomotor activity, (B) proportion of time flies spent sleeping and (C) mean activity while flies were awake, during the first 96 hours of DCV infection. Across all panels, sex and infection status are represented by colour with uninfected females shown in blue, infected females in pale blue, uninfected males in red, and infected males in pink. The order of genetic backgrounds on the x-axis of each of panel follows the ascending order of female flies.

## Discussion

Identifying changes in host behaviour following infection is important to understand heterogeneity in disease transmission. Overall, our results indicate a significant sex difference in the effect of infection on social aggregation but no effect of infection on locomotor activity in either sex.

We observed that how closely flies aggregate with one another differs with their genetic background. The genetic variation we observed is similar to other studies that have measured nearest neighbour distance [10], as well as other aspects of *Drosophila* social behaviour, such as group size preference [9] and group composition [28]. Group composition is affected by the natural *foraging* gene polymorphism, where larvae are either sitters, which aggregate toward conspecifics at food sources or rovers, who are more prone to independent food searching behaviour. Larger groups of larvae on food patches are more likely to be comprised of sitters, as rovers leave food patches after overcrowding [28]. Genetic components of social behaviour have also been identified in a number of mammal species, including humans [29]. In a number of vole species, variation in oxytocin [30] and arginine vasopressin [31] receptor density is associated with between-species variation in pair-bonding and monogamy.

While healthy male and female aggregation did not differ, once infected, males aggregated further apart, while female aggregation did not change. One possible explanation for why males aggregate further apart following infection is a sex difference in immunity and the costs of social aggregation [32]. Sexually dimorphic immunity may be particularly relevant as male *D. melanogaster* exhibit a stereotyped suite of aggressive behaviours [33–35]. While fighting can gain males access to valuable resources, it often incurs substantial costs [36,37]. DCV infection could exacerbate the cost of male aggregation, as resources would also need to be spent on fighting infection, which could lead to males aggregating less. Despite females also fighting one another, this aggression is generally less costly [38,39]. Females may therefore still be able to aggregate relatively closely while fighting DCV infection.

Irrespective of the metric used, we found no measurable effect of DCV infection on locomotor activity. Other work has shown decreases in *Drosophila* daily movement following injection with DCV, where a marked reduction is seen after 4 days of infection [40]. Reduced daily locomotor activity has also been observed in *Drosophila* after 3 days of infection with the DNA virus Kalithea virus [41]. Injecting, rather than pricking, flies with viral suspension, allows more precise control of infectious dosage, which could also increase infection severity [42]. Another potential explanation is that we infected flies via thoracic pricking, as opposed to abdominal injection which has been shown to reduce resistance to infection in *Drosophila* [43]. Orally infecting flies shows a range of sex-specific behavioural symptoms, with sublethal doses reducing daily locomotor activity in males after 3-6 days of infection [24]. Conversely, following oral infection with larger doses of DCV, females, but not males, have been shown to sleep more [23]. These studies suggest we may not have seen an effect of DCV infection on activity, because infections were not severe enough to elicit behavioural symptoms or that any effect was concealed by a response to thoracic injury. Measuring the activity of flies later in infection might address these explanations, as this will enable flies to heal from thoracic injury and accrue a greater viral burden.

We measured social aggregation in groups of individuals composed of the same genetic background, sex and infection status in order to dissect their influence on social aggregation. However, in more heterogenous wild populations these characteristics can produce population structure that could affect contact between individuals. Individuals with shared genotypes can be more likely to interact due predispositions to traits such as group size preference [28,44] and aggression [45]. Similarly, sexual interactions between males and females, as well as fighting and other forms of sexual competition, further alter contact networks within populations [46,47]. When present together, healthy hosts might also be able to avoid infected conspecifics by detecting the pathogen or cues of its pathology [48]. Future work aiming to characterise the influence of these sources of variation on heterogeneity in contact rate should consider how they change with, and are changed by, population structure.

The contrasting ways social aggregation and locomotor activity change following infection highlight the complexity of sources determining between-individual variation in disease transmission. This is complicated further by sex differences across and within these genetic backgrounds. The change induced by DCV infection on social aggregation but not locomotor activity also demonstrates the importance of considering multiple host behaviours. Central to understanding the effect of this genetic and sex-specific variation in social aggregation and locomotor activity on heterogeneity in disease transmission is characterising their effect on contact rates. Additionally, future work should consider how these traits interact with other key determinants of transmission, such as infectiousness and infection duration, as these three components ultimately define disease transmission in conjunction with one another.

## Acknowledgements

We are grateful to Katy Monteith for general lab assistance, Angela Reid and Lucinda Rowe for preparation of all fly media and the Regan, Obbard, Walling and Vale lab members for constructive discussion of these ideas.

## Accessibility

All data and R code will be uploaded to Dryad during submission.

## Funding

J.A.S.J. was supported by a NERC E3 DTP PhD studentship. P.F.V. was supported by a Branco Weiss fellowship (https://brancoweissfellowship.org/) and a Chancellor’s Fellowship (School of Biological Sciences, University of Edinburgh).

### Ethical Statement

This work poses no ethical concerns.

## Electronic Supplementary Material

### Detailed Materials & Methods

#### Social aggregation experiment

Social aggregation was measured in 55mm Petri dishes with 2% agar poured in until 3mm from the lid in order to limit flight. Flies were pricked with DCV or TRIS as described in the main methods and, under light CO_2_ anaesthesia, transferred to Petri dishes in groups of twelve. Due to reducing anaesthesia as much as possible to curtail behavioural defects associated with over-exposure to CO_2_ [1], and experimenter error, some flies escaped Petri dishes before they were closed. A total of 448 dishes contained twelve flies, while 113 and 19 contained eleven and ten, respectively. Flies within a Petri dish were the same genetic background, sex and infection treatment. Once transferred, flies were left in Petri dishes to acclimate for 30 minutes. This acclimation period was identified in a prior experiment where it was observed that after 30 minutes, fly movement in arenas was minimal, as shown previously [2,3]. A single image was recorded of each Petri dish using a 13 Megapixel camera, followed by a second image (10-20 minutes later). Using these images we calculated the NND using ImageJ software [4], by marking flies in the centre of their thorax with the multi-point tool. We calibrated the distance between flies in photos using the 55mm width of the Petri dish and calculated the nearest neighbour distance between each pair of flies in millimetres using the ‘NND’ package in ImageJ. These values were used to calculate the median NND for each petri dish [2,5]. To account for differences in body lengths between lines and sexes, we also calculated the NND using body lengths by dividing millimetre distances by the mean body length of a randomly selected group of 30-40 individuals from each genetic background and sex combination (Figure S1). Incorporating this size difference into measures of social aggregation, by measuring body lengths between individuals did not alter the results qualitatively (Figure S2).

**Figure S1.**
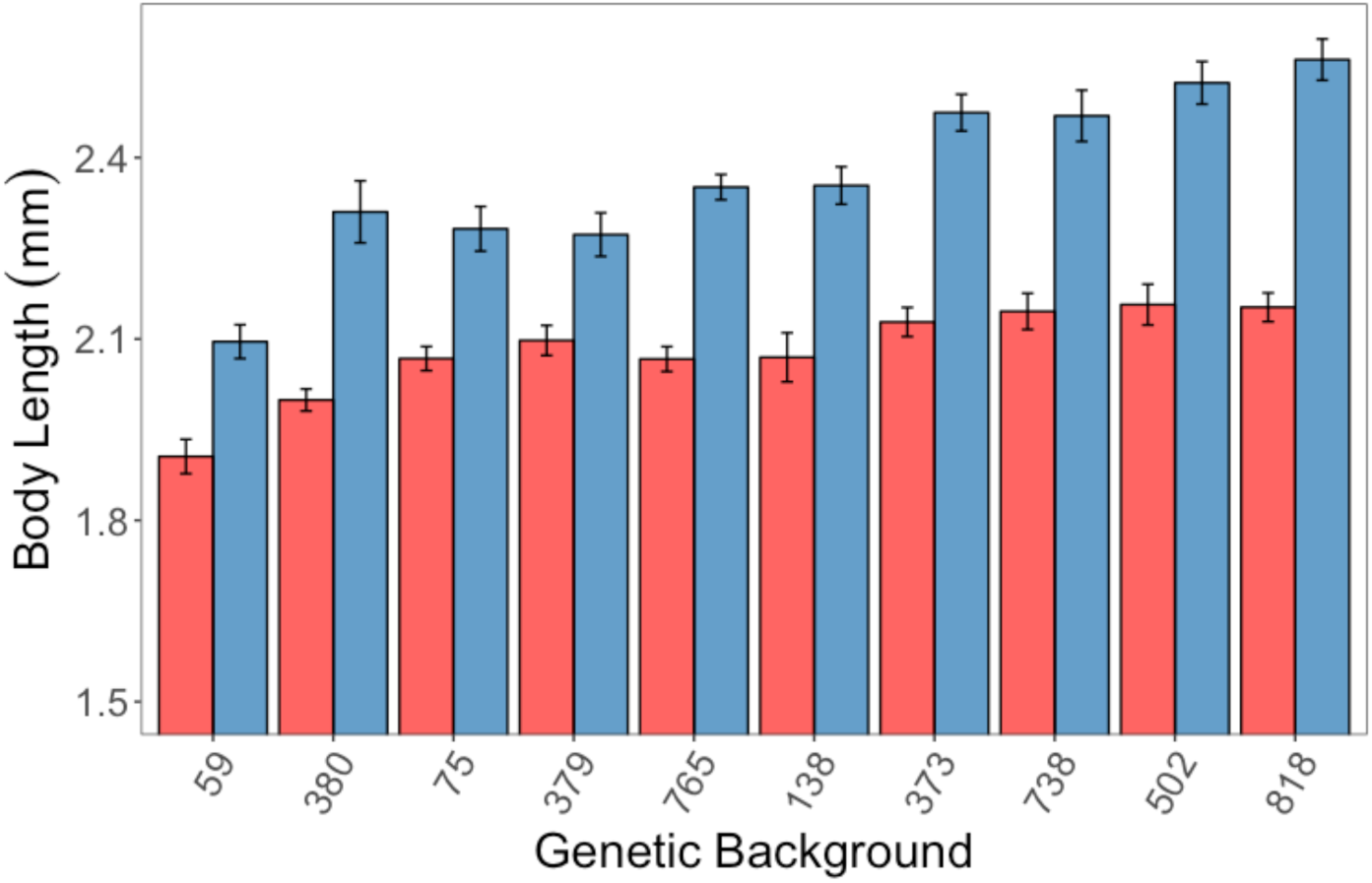
Mean±SE body length of flies calculated from 30 flies per line per sex.

**Table S1.**
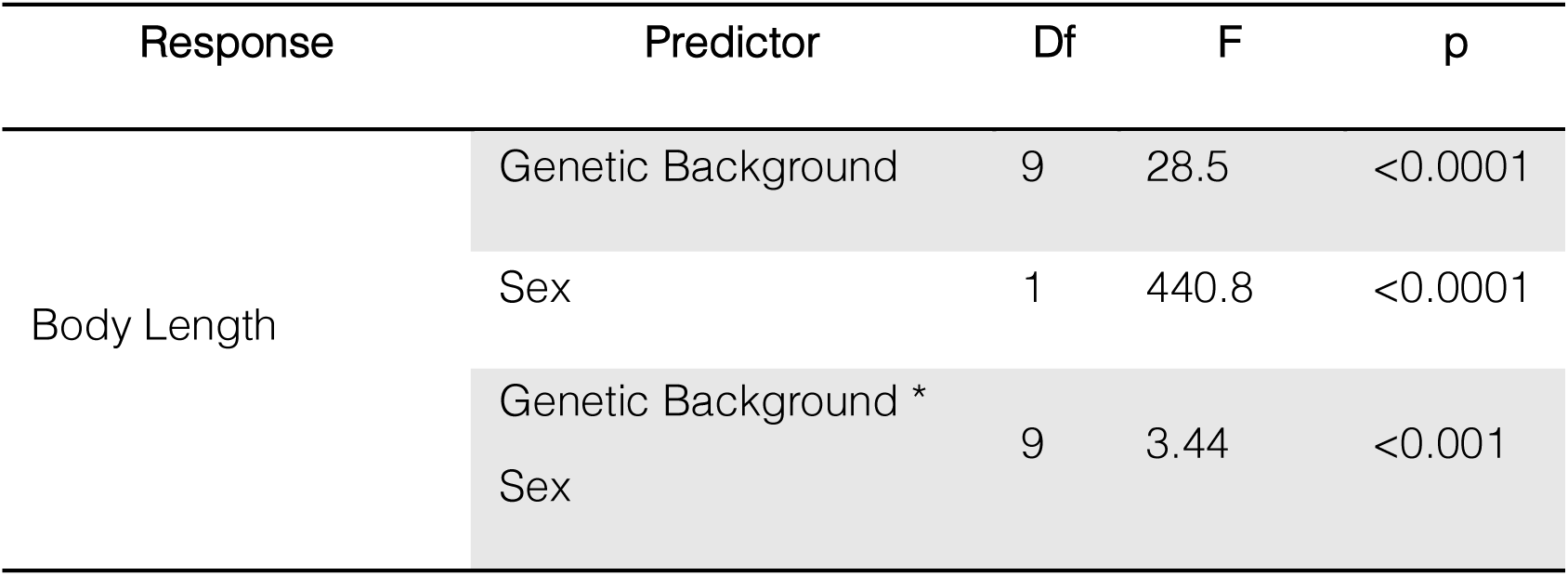
Model outputs for statistical tests performed on body lengths of treatment groups comprised of each combination of sex and genetic background.

**Figure S2.**
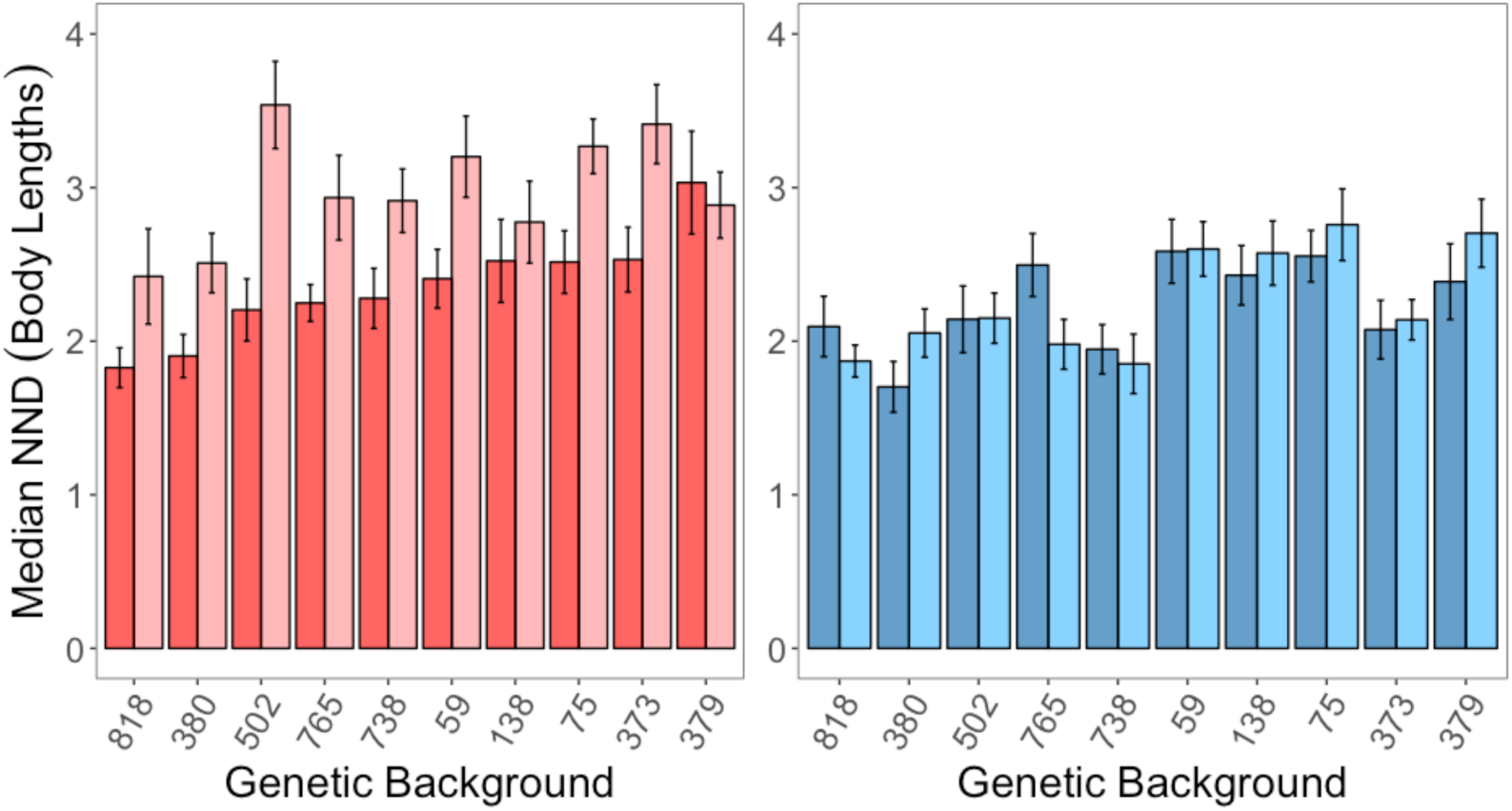
Mean±SE median nearest neighbour distance (NND) in body lengths of adult flies placed in Petri dishes for at least 30 minutes until settled. (a) Uninfected female-only arenas shown in blue, and infected female-only arenas in pale blue. (b) Uninfected male-only arenas are shown in red, and infected male-only arenas in pink. The x-axis of both panels is ordered from the lowest to highest mean median NND of female flies of a single genetic background.

**Table S2.**
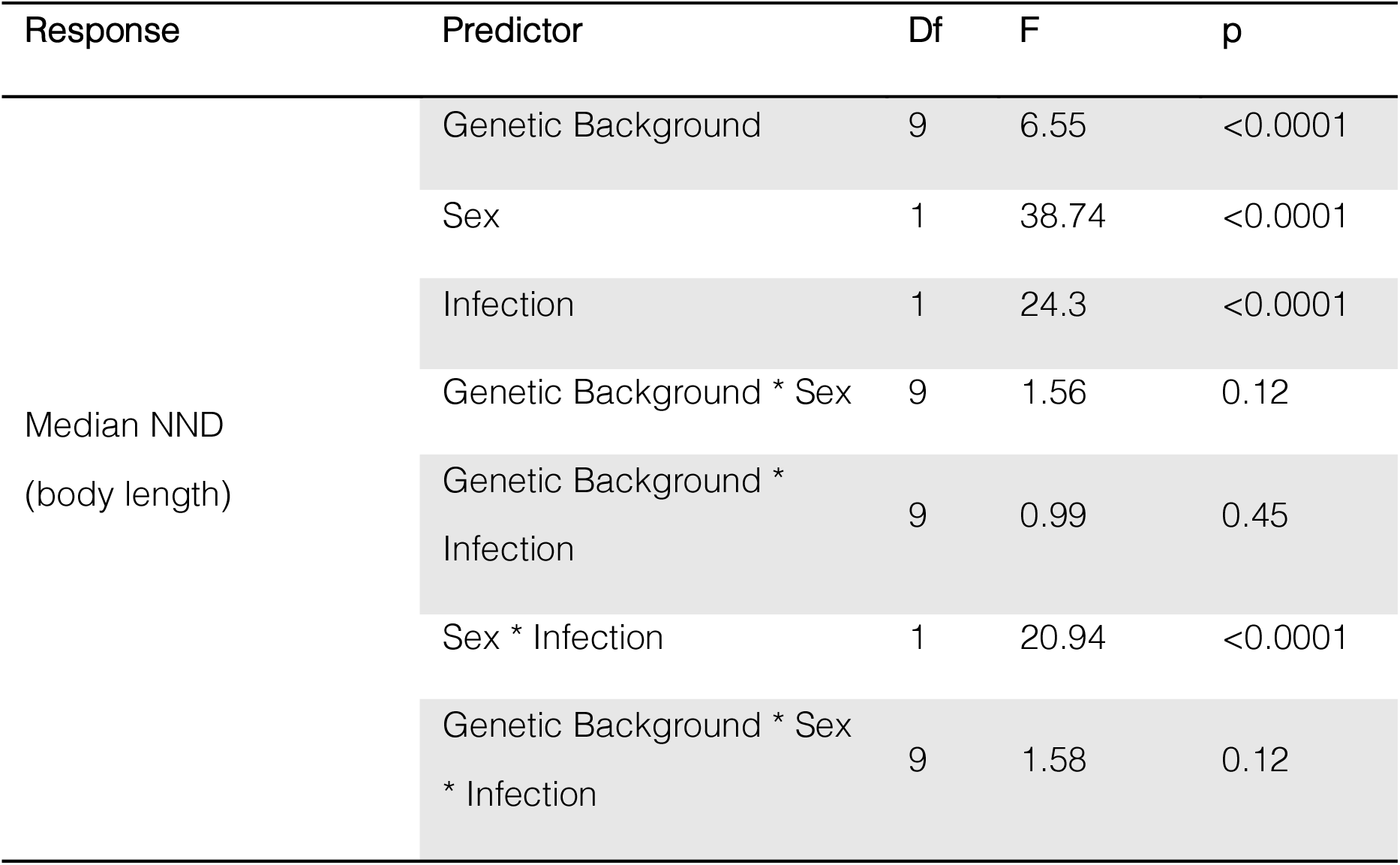
Model outputs for statistical tests performed on social aggregation when measured using body lengths, testing the causes of variation in sociality in males and females of 10 *D. melanogaster* genetic backgrounds.

| Response | Predictor | Df | F | p |
| --- | --- | --- | --- | --- |
| Median NND<br>(body length) | Genetic Background | 9 | 6.55 | <0.0001 |
|  | Sex | 1 | 38.74 | <0.0001 |
|  | Infection | 1 | 24.3 | <0.0001 |
|  | Genetic Background * Sex | 9 | 1.56 | 0.12 |
|  | Genetic Background *<br>Infection | 9 | 0.99 | 0.45 |
|  | Sex * Infection | 1 | 20.94 | <0.0001 |
|  | Genetic Background * Sex<br>* Infection | 9 | 1.58 | 0.12 |

#### Analysis of Drosophila Activity

Single flies were either systemically infected or control pricked as described above, and immediately placed in a single DAM tube and allocated a random slot in one of 8 DAM monitor units (each unit is capable of housing a maximum of 32 tubes). At least one slot of each monitor unit was left empty and another contained an empty tube, as negative controls. While flies were monitored continuously for 4 complete days. Flies that died during this 4-day period were removed from the dataset. In total we analysed the activity of 872 flies, with n=18-27 flies for each combination of sex and genetic background (Table S3). Raw activity data was processed using the DAM System File Scan Software [6], and the resulting data was manipulated using Microsoft Excel. Activity counts for each individual fly were combined into 5-minute bins. We analysed fly activity data using three metrics: total locomotor activity, proportion of time spent asleep and the average activity when awake [7]. Total locomotor activity refers to the sum of all recorded movements during the 4-day measuring period and is an outcome of how often a fly sleeps and how much it moves during bouts of awake activity. In Drosophila, sleep is defined as five minutes of continuous inactivity, sharing several features with mammalian sleep, such as being followed by an increased arousal threshold, and being regulated independently from the circadian clock [8]. To assess the proportion of time spent asleep, we used the proportion of all 5-min bouts (n=1152) where no activity was logged. To quantify awake activity, we calculated the average level of locomotor activity across every 5-min period where at least one instance of movement was recorded. Average activity when awake can help characterise lethargy when individuals are active, an important behavioural symptom of infection [9].

**Table S3.** The sample size of each activity treatment group, representing every combination of sex, genetic background and infection status.

| Genetic<br>Background | Male |  | Female |  |
| --- | --- | --- | --- | --- |
|  | Control | Infected | Control | Infected |
| RAL-59 | 24 | 24 | 22 | 22 |
| RAL-75 | 20 | 27 | 20 | 20 |
| RAL-138 | 21 | 20 | 18 | 19 |
| RAL-373 | 20 | 26 | 21 | 28 |
| RAL-379 | 20 | 22 | 21 | 20 |
| RAL-380 | 24 | 20 | 23 | 24 |
| RAL-502 | 23 | 21 | 21 | 20 |
| RAL-738 | 21 | 21 | 20 | 20 |
| RAL-765 | 26 | 28 | 20 | 21 |
| RAL-818 | 21 | 20 | 20 | 22 |

**Figure S3.**
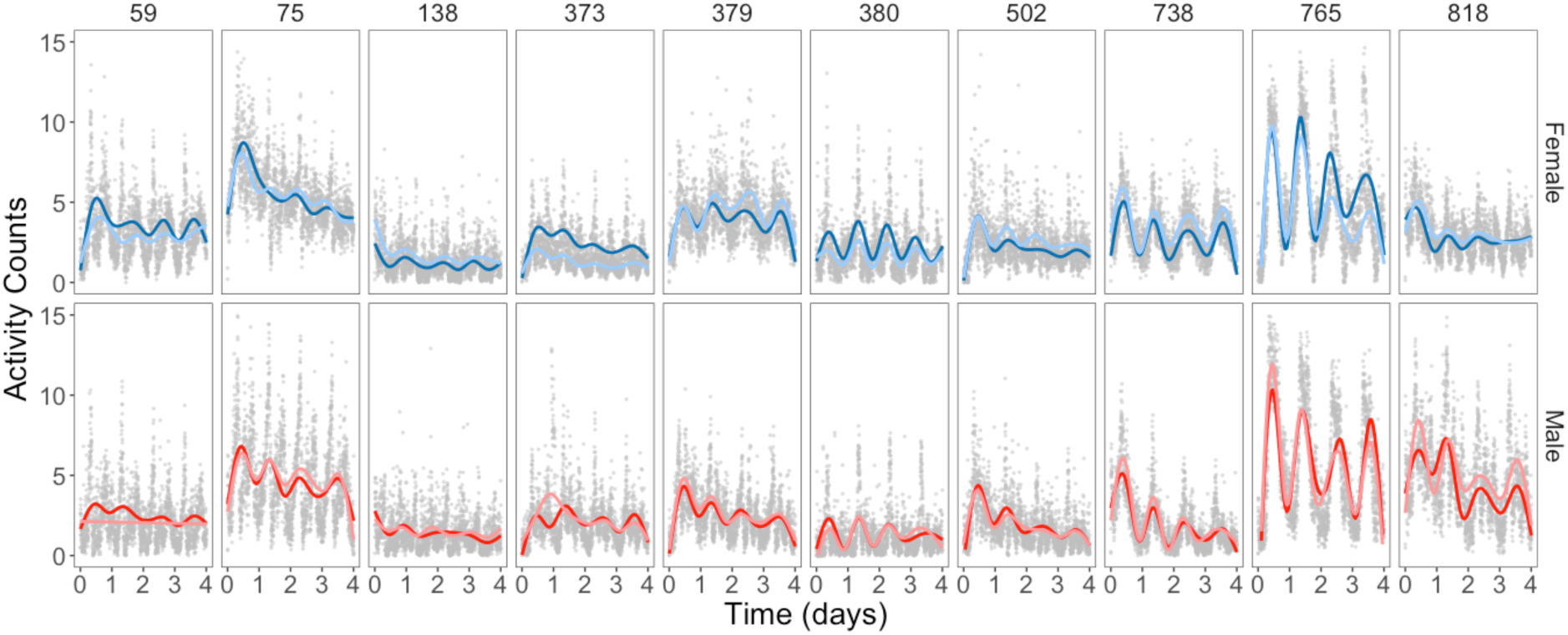
Activity counts of adult flies for the first 4 days locomotor activity was measured in the DAM. The mean activity counts of DAM tubes containing single flies of the same sex and DCV infection status are represented by generalised additive model lines where uninfected females are shown in blue, infected females in pale blue, uninfected males are shown in red, and infected males in pink.

